# Insulin evokes release of endozepines from astrocytes of the NTS to modulate glucose metabolism

**DOI:** 10.1101/2024.05.03.592384

**Authors:** Lauryn E. New, Niannian Wang, Shabbir Khan, Joanne C. Griffiths, Ryan Hains, Jamie Johnston, Beatrice M. Filippi

## Abstract

The central nervous system (CNS) plays a key role in regulating metabolic functions, but conditions like obesity and diabetes can disrupt this balance. Within the CNS, the nucleus of the solitary tract (NTS) in the dorsal vagal complex (DVC) controls glucose metabolism and feeding behaviour. In rodents, the NTS senses insulin and communicates with the liver to regulate glucose production. Even short-term exposure to a high-fat diet (HFD) can lead to insulin resistance and impair NTS function. However, we still know little about which cells in the NTS are sensitive to insulin. Our study aimed to identify these insulin-sensitive cells and understand how they affect glucose metabolism.

We found that insulin receptors in astrocytes are crucial for the NTS’s ability to regulate glucose production in the liver. Insulin evokes the release of endozepines from astrocytes, and injecting endozepines into the NTS reduces glucose production. The effect of endozepines within the NTS is mimicked by GABAA antagonists and prevented by an agonist, suggesting that insulin prompts astrocytes to release endozepines, which then attenuate GABAA receptor activity, ultimately reducing glucose production in the liver.

Our study is the first to show that insulin–dependent release of endozepines from NTS- astrocytes is fundamental to control blood glucose levels, providing valuable insights into the mechanisms underlying insulin function within this specific region of the CNS.

## Introduction

The maintenance of glucose homeostasis requires the fine balance between the release and uptake of glucose by different organs, all coordinated by the modulation of insulin and glucagon levels. The interaction between pancreatic islets and insulin- sensitive tissues has been traditionally used to explain how glucose homeostasis is maintained in the body, but compelling evidence for an important role for the central nervous system (CNS) continues to accumulate^1^. The CNS can sense the metabolic state of the body via sensory inputs from the vagus nerve and by directly sensing hormones and nutrients and feeds back to peripheral organs to maintain energy balance. Hormones like insulin, leptin, GLP-1, and ghrelin act on brain receptors to regulate metabolic balance by activating/inhibiting specific neuronal populations to maintain metabolic homeostasis^2–5^. Insulin can pass the blood brain barrier and reach different brain areas to lower plasma glucose levels ^6,7^ and this is in part explained by a decrease in hepatic glucose production (HGP) ^8^, which is communicated by the vagal nerve ^9^. Several brain regions have been identified that sense insulin and regulate HGP, including the arcuate nucleus of the hypothalamus ^9^ and more recently the Nucleus of the Solitary Tract (NTS) in the Dorsal Vagal Complex (DVC) ^10,11^. Insulin modulates neural excitability in these brain regions by activating the ATP-sensitive potassium channel (KATP) ^10,11^; in the arcuate nucleus it results in inhibition of both POMC and AGRP neurons^11^, whereas the neural targets of insulin with the NTS have not yet been identified.

The NTS receives sensory information from multiple types of vagal afferent fibres that carry information from the viscera. The NTS then integrates and relays this information to the hypothalamus and the dorsal motor nucleus of the vagus (DMX). The hypothalamus regulates food intake and metabolic behaviour, while the DMX signals back to the viscera to coordinate gastric reflexes, motility, emptying and to the liver to modulate HGP ^12,13^. The NTS contain neurons that release and are sensitive to a large array of neurotransmitters, including γ-aminobutyric acid (GABA) ^14^, glutamate ^15^ and catecholamines ^16^, as well as neuropeptides such as glucagon-like peptide 1 (GLP-1)^17^ and neuropeptide Y (NPY) ^5^. An essential yet unanswered question is identifying the specific NTS cell types involved in regulating food intake and hepatic glucose HGP in response to insulin. Interestingly, astrocytes regulate many aspects of neuronal function, including synaptic plasticity, survival, metabolism, and neurotransmission^18^. Chemogenetic activation or inhibition of NTS astrocytes affects feeding behaviour^19^ and inhibiting mitochondrial fission in NTS astrocytes protects rats from high-fat diet-induced insulin resistance and obesity^20^. Intriguingly, specific knockdown of the insulin receptor in astrocytes of the mediobasal hypothalamus (MBH), causes hyperphagia and affects glucose metabolism and energy expenditure ^21,22^. Therefore, astrocytes are likely to be important for insulin sensing within the NTS but the specific details have not yet been determined. One mechanism of astrocyte-neuronal cross talk involves the release of astrocyte specific peptides such as endozepines, notably the 86-amino acid precursor diazepam-binding inhibitor (DBI) and its proteolytic cleavage product, octadecaneuropeptide (ODN) which act on GABAA receptors to modulate neuronal activity ^23–25^. DBI and its derived peptides have been shown to play a central role in systemic metabolic regulation^26^.

Here we investigated the possibility that insulin could act on astrocytes in the NTS to decrease HGP and control blood glucose levels. We discovered that insulin can trigger the release of endozepines from astrocytes which modulates GABAergic signalling in the NTS, resulting in decreased hepatic glucose production.

## Results

### INSR is preferentially expressed in astrocytes and VGAT neurones of the NTS

The NTS is comprised of a plethora of different neuronal cell types including glutamatergic, GABAergic, and dopaminergic neurones among others^27,28^. The lack of information regarding the distribution of the insulin reception in the NTS prompted us to perform RNAScope focusing on 3 major cell types; astrocytes, GABAergic and glutamatergic neurones (Fig. 1 A and B). We found that a large fraction of cells within the NTS were positive for the insulin receptor, 64 ± 2.1% of DAPI identified cells. Of these insulin positive cells, 38 ± 2.1 % were GABAergic with only 16 ± 1.5 % being glutamatergic; we also found that GABAergic neurons outnumbered glutamatergic by ∼3:1 (Fig. 1D). However, the largest population of insulin positive cells were astrocytes (Figure 1 B&C). Overall, these data suggest that when insulin is elevated multiple cell types within the NTS may be modulated at the same time.

**Figure 1:**
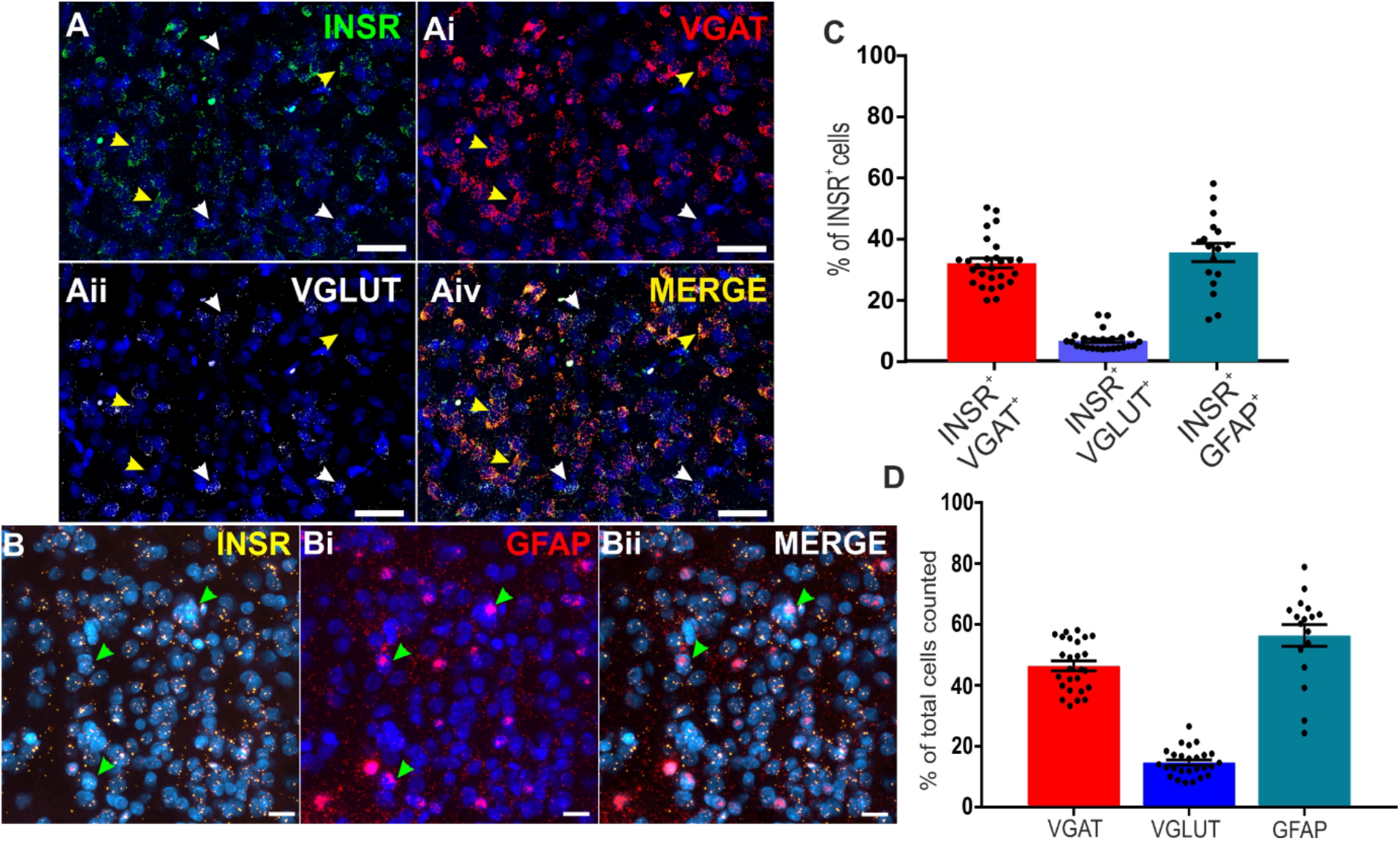
Neuronal population and insulin receptor distribution in the NTS: **(A and B)** RNAScope staining of the NTS area showing the localization of the insulin receptor (INSR, **A** Green and **B**, Yellow) with GABAergic neurones (VGAT, **Ai**, Red), glutamatergic neurones (VGLUT, **Aii**, White) and Astrocytes (GFAP, **Bi**, Red). Merged images are shown in **Aiii** and **Bii**. A-AiV Scale bar=50μm, B-Bii scale bar = 20μm (3 ROIs counted from 3 sections per animal, N= 3 animals). **(C)** The % distribution of INSR+ cells which were also VGAT^+^, VGLUT^+^, or GFAP^+^ in the NTS. **(D)** The % distribution of astrocytes, GABAergic neurones and glutamatergic neurones in the NTS.

### Insulin induces widespread c-Fos expression in the NTS, predominantly in cells lacking insulin receptors

We have shown that insulin receptors are expressed by multiple cell types within the NTS (Fig. 1), we next sought to address how insulin modulates activity within the NTS. We injected FITC-labelled insulin (F-ins) into the NTS through a brain cannula (supplementary figure S1) and then stained for the immediate early gene c-Fos, which gets induced by increased neural activity^29^. FITC-insulin acts like normal insulin, triggering AKT activation in PC12 cells (supplementary figure S2) and importantly it gets internalised along with the activated receptor and therefore accumulates inside the cell^30^ enabling us to identify cells in which the insulin receptor had been activated. We found a striking increase in c-Fos in NTS that were injected with F-ins compared to the contralateral side injected with saline (822.2 ± 84.1 vs 176.4 ± 15.9, Figure 2 A-D, p<0.0001). This indicates that elevated insulin markedly increases activity within the NTS, however the canonical mode of action is for insulin to activate the KATP channel, which would reduce neural activity^9,10^. Indeed, we found that the majority of FICT-insulin labelled cells were not c-FOS positive, indicating that the increased c-FOS expression observed in Fig. 2 A-D is likely due to an indirect action of insulin. This may be due to insulin decreasing activity in GABAergic neurons (Fig. 1C), leading to increased activity of other neurons due to disinhibition. Alternatively, the 22% of cells that were positive for FITC-insulin and c-FOS (Fig. 2E) and therefore directly activated, may be exerting a net excitatory effect on a larger population of neurons.

**Figure 2:**
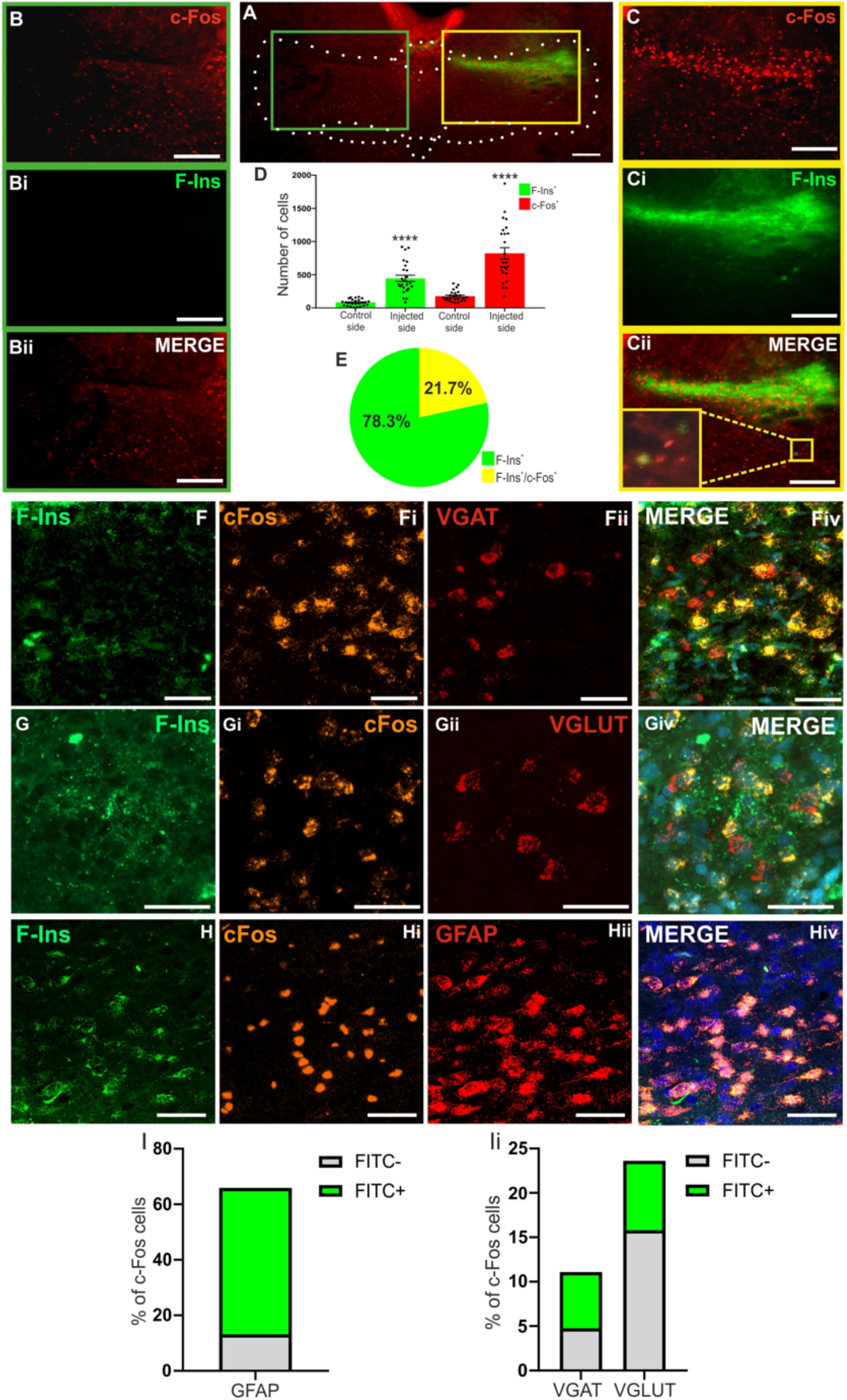
F-Ins injection increases c-Fos levels in the DVC. (A-Cii) Representative slidescanner images showing c-Fos (red) and F-Ins (green) staining, and c-Fos and F-Ins merged images in non-injected (green box, B-Bii) and injected DVC areas. (yellow box, C- Cii) Closed arrows denote colocalised cells. Open arrows denote non-colocalised cells. Scale bar=50µm. **(D)** Comparison of cells counts of F-Ins-positive and c-Fos-positive cells overall in DVC sections in animals injected or non-injected with F-Ins. (*N*=2 *n*=25, where N is the number of animals and n is the number of sections counted). **(E)** Percentage of total F-Ins+ cells which were either F-Ins positive only (green wedge) or F-Ins^+^/C-Fos^+^ (yellow wedge). **(F-Hiv)** RNAScope staining of the NTS area looking at the localization of FITC-insulin (F**, G and H)** and *C-Fos **(F***i, G**i** and H**i**) with *VGAT* (F**ii**), *VGLUT* (G**ii**) or *GFAP* (H**ii**). Merged images are shown in Fiv **and Giv and Hiv**. Scale bar=50µm. (I-Ii) The % of c-FOS+ cells which are negative for FITC-insulin (grey bar) or positive for FITC-insulin (green bar) which colocalise with probes for GFAP (I) The % of c-FOS+ cells which are either FITC-insulin^-^ (grey bars) or FITC-insulin^+^, that co-localise with probes for either VGAT or VGLUT (3 ROIs counted from 3 sections per animal (*N*=3) where *n*=27 where *n* is the number of ROIs quantified per animal)

To determine which cell types in the NTS become activated with insulin application we combined RNAscope with the FITC-insulin injection (Fig 2F-hiv). The largest fraction of FITC-insulin and c-FOS positive cells we identified as astrocytes (52.8 ± 3% of c- Fos+ cells, Fig 2I), whereas only 6.3 ± 0.8% and 7.8 ± 0.92 % of c-Fos+/FITC+ cells were VGAT+ or VGLUT+, respectively (Fig 2Ii). Together these data show that insulin causes broad activation of NTS cell types and that astrocytes appear to be directly activated by insulin, whereas other neuronal cell types are activated via a predominantly indirect means, possibly involving activation of astrocytes.

### Knockdown of insulin receptors in NTS-astrocytes impaired insulin-dependent decrease of hepatic glucose production

We have identified that insulin is likely to directly activate astrocytes within the NTS, so we next determined whether astrocytic insulin sensing is necessary for the NTS control of the brain-liver axes that modulate blood glucose levels. To this aim we developed a Cre-lox system to specifically knock down the insulin receptor in astrocytes. We first validated this system in vitro by co-expressing the Cre recombinase together with a FLEx plasmid where ShRNA for the insulin receptor and tdTomato (ShIR) are cloned within 2 LoxP sites. Compared with scramble (ShC) expressing cells, the expression of ShIR decreased insulin receptor levels by 40% (Figure 3A). Insulin ShIR expressing cells were not able to uptake FITC-insulin when compared with ShC expressing cells or their untransfected neighbours (Figure 3 B to D), which indicates that the insulin receptor is required for FITC-insulin labelling of cells in Fig. 2. We then developed 2 different adenoviruses, one expressing the Cre recombinase -CFP tagged under the GFAP promoter (GFAP-Cre) to specifically target GFAP-expressing astrocytes and another one expressing the ShRNA for the insulin receptor and tdTomato under 2 lox P sites. Primary astrocytes incubated with these 2 viruses showed a 60% reduction of the insulin receptor levels and impaired insulin signalling, as shown by decreased AKT phosphorylation (Figure 3E).

**Figure 3.**
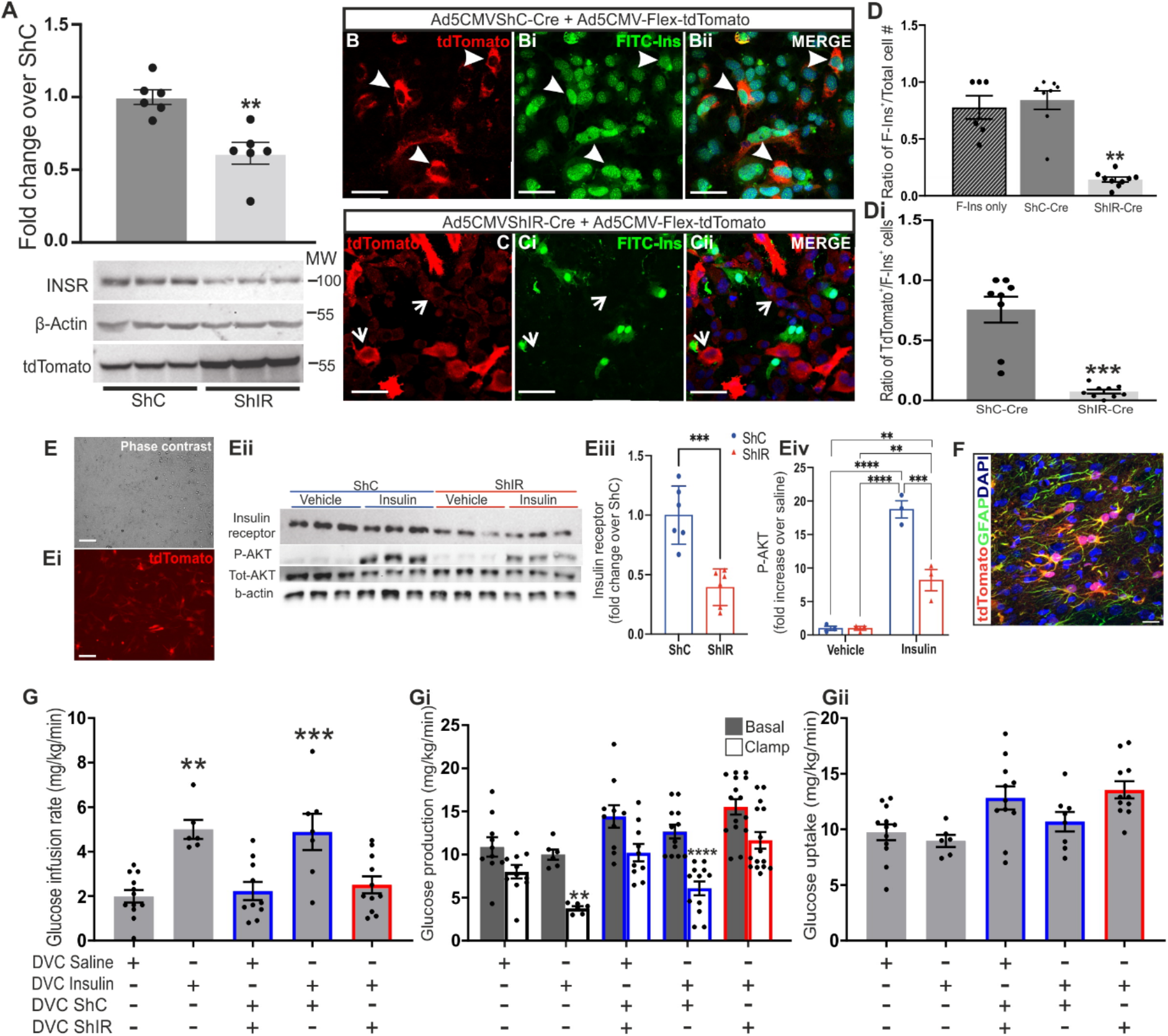
Effect of the knock down of insulin receptor in NTS GFAP+ astrocytes on insulin- dependent regulation of glucose production. (**A-D**) PC12 cells co-transfected with a Cre expressing plasmid and a FLExed plasmid expressing the ShRNA for the insulin receptor (ShIR) or a scramble ShRNA (ShC). (**A**) Quantification of insulin receptor levels and tdTomato’s levels. (**B-D**) Cells were starved overnight and then treated with 800nM FITC- insulin. Representative image of FITC-insulin binding is shown in B and C and quantification of FITC- insulin binding is shown in D and Di. Closed arrows indicate co-localization between tdTomato and FITC-insulin, closed arrows indicate tdTomato-positive cells without FITC-insulin. (**E**) Primary astrocytes where infected with 2 adenoviruses expressing Cre-recombinase and a FLEx virus expressing the ShIR or the ShC. Cells where then starved for 5 hours and treated with 100nM insulin or saline for 30 min. Cells where then harvested lysed and analysed by western blotting and probed with the indicated antibodies. The figure shows a representative image of the primary astrocytes expressing tdTomato (E) and a western blot with insulin receptor and p-AKT quantification normalised for b-actin and total-AKT respectively (Eii-Eiv). (**F**) Following clamp animals were fixed 4% with PFA and 25 µm coronal sections of the DVC areas were stained with GFAP to visualise astrocytes. tdTomato shows cells expressing the ShRNA for the insulin receptor (ShIR). Nuclei are stained with DAPI. Arrows indicates co-localization between td-Tomato and S100B. Scale bar = 20µm (see also supplementary figure S3 for single channel images). (**G-Gii**) The day of brain surgery rats were co-injected with an adenovirus expressing the Cre recombinase under the GFAP promoter (GFAP-Cre) and with an adenovirus expressing either the ShIR or the ShC. Saline or insulin (a total of 2.52 mU, 1.26 mU per site) were infused at 0.006 ml/min during the clamp through the DVC catheter (see also Supplementary figure S4). Basal glucose levels measured in food restricted rats the morning of the clamp (G). Glucose infusion rate (mg/Kg/min) during the final 30 min of the clamp for the indicated treatment (Gi). Glucose production (mg/Kg/min) in basal (with square) and clamp (black square) conditions (Giii) Glucose uptake (mg/Kg/min) during the final 30 min of the clamp for the indicated treatment. Values are shown as mean + SEM; *n*= 12 for saline, *n*= 6 for insulin, *n*= 11 for ShC saline, (mix of ShIR and ShC rats), *n*= 7 insulin ShC, *n*= 11 insulin ShIR

Rats that received an NTS-targeted co-injection of the 2 adenoviruses (2.5ul each side) expressed the Cre recombinase only in astrocytes, leading to the specific expression of tdTomato and ShIR in this cell type (co-localization between tdTomato and GFAP, Figure 3F). We then performed a pancreatic euglycemic-basal insulin clamp study^10^ where rats were injected either with insulin or saline in the NTS and hepatic glucose production (HGP) was measured with the tracer dilution methodology (supplementary figure S4). Somatostatin was used to prevent changes in circulating glucoregulatory hormones such as insulin, which were experimentally maintained at basal level (Supplementary Table S1), while tritiated glucose was infused intravenously to measure the amount of glucose that is taken up by peripheral organs or released by the liver (GU and HGP respectively, supplementary figure S4). The clamp was performed in rats restricted to 70% of their average food intake overnight. In control (ShC rats), upon insulin injection in the NTS, we could clearly see an increase in the amount of infused glucose required in order to keep glycemia at basal level (glucose infusion rate, GIR, Figure 3 G, supplementary table S2). The increase in GIR suggested that NTS infusion of insulin was decreasing blood glucose levels. To determine whether this decrease was due to an increased GU or a decreased HGP, we applied tracer dilution methodology^31^. The higher GIR was associated with a decrease in HGP while the glucose uptake was unchanged (as previously reported in Filippi et al, (2012), Figure 3 Gi and Gii). NTS-insulin in ShC animal was able suppress HGP by 60% (supplementary figure S5). However, expression of ShIR to specifically knockdown the insulin receptor in astrocytes was sufficient to block insulin action thus confirming that insulin acts through astrocytes in the NTS to decrease HGP (Figure 3G to Giii and S5).

### Endozepines are released by astrocytes upon insulin treatment and can decrease HGP

We have shown that insulin can directly activate astrocytes within the NTS (Fig. 2) and that insulin sensing by astrocytes is required for the NTS to respond to insulin by decreasing HGP (Fig. 3). How could NTS astrocytes modulate the activity of NTS neurons to enable communication to the liver? The endozepine DBI is enriched in astrocytes^25^, this molecule and its metabolite ODN can be released from glia and modulate the function of GABAA receptors. We therefore tested whether insulin evokes release of DBI from astrocytes. Insulin treatment of primary astrocyte cultures increased the phosphorylation levels of AKT (Figure 4 Ai) and ERK1/2 (Figure 4 Aii) as expected. Strikingly, there was also 40% increase in the level of DBI in the media as measured with an ELISA assay (Fig 4Aiii) indicating that insulin triggers the release of DBI from primary astrocytes.

**Figure 4:**
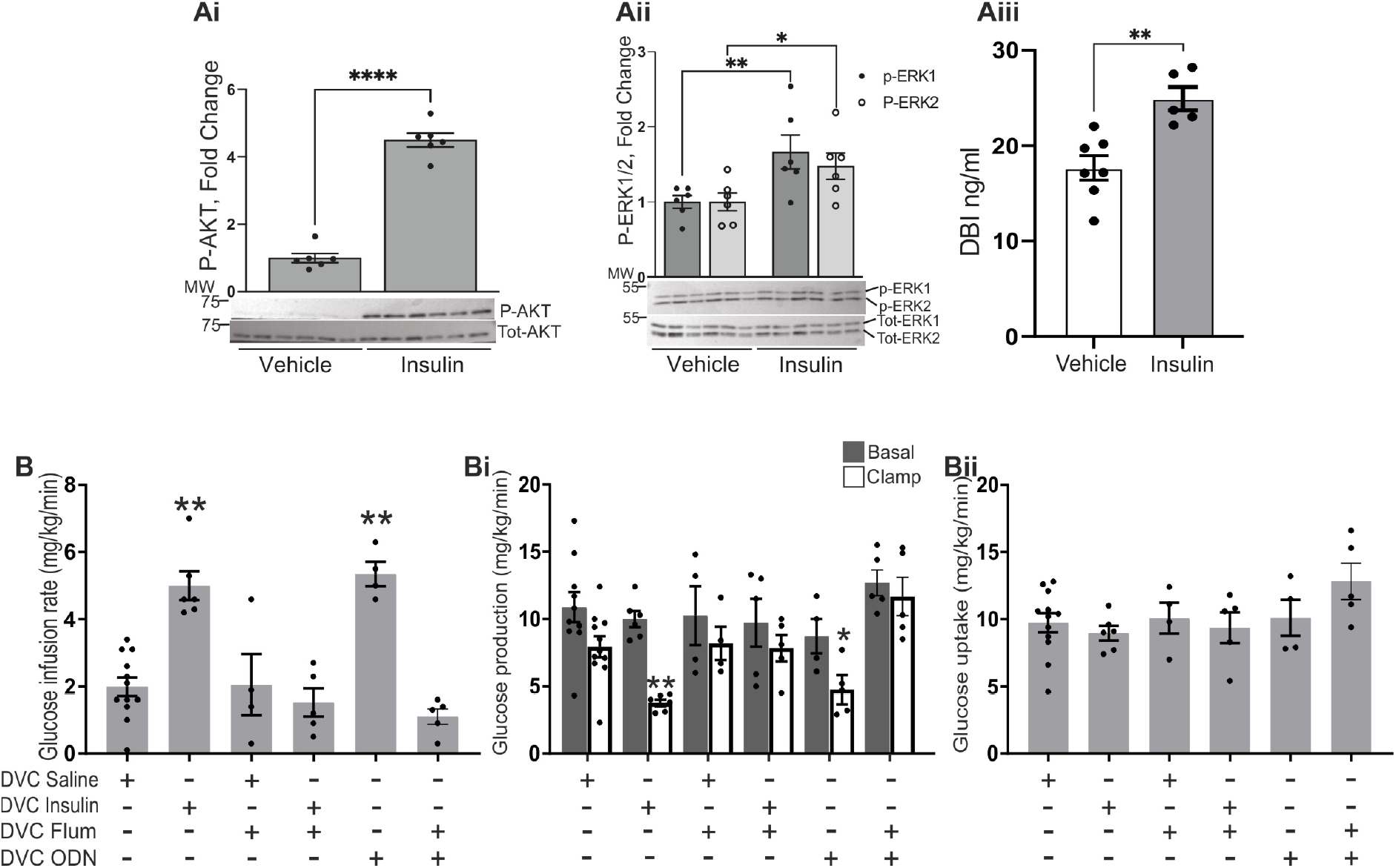
Endozepines are released by astrocytes upon insulin treatment and can decrease HGP. (Ai-Aii) Primary astrocytes were starved for 5 hours and treated with 100nM insulin or saline for 30 min. Cells where then harvested lysed and analysed by western blotting and probed with the indicated antibodies. The figure shows a representative western blot with Insulin receptor and p-AKT quantification normalised for b-actin. (Aii) A comparison of the level of DBI (ng/ml) present in the media from the primary astrocyte experiment shown in Ai-Aii as measured by ELISA. (B-Bii) Data from pancreatic euglycemic clamp experiments where either saline, insulin (a total of 2.52 mU, 1.26 mU per site), flumazenil ([33nM]), ODN ([10mM]) or flumazenil ([33nM]) plus ODN ([10mM]) were infused at 0.006 ml/min during the clamp through the DVC catheter. (**B**) Glucose infusion rate (mg/Kg/min) during the final 30 min of the clamp for the indicated treatment. (**Bi**) Glucose production (mg/Kg/min) in basal (with square) and clamp (black square) conditions. (**Bii**) Glucose uptake was comparable in all groups. Values are shown as mean + SEM; *n*= 12 for saline, *n*= 5 insulin, *n*= 4 for flumazenil and 5 for flumazenil plus insulin, *n*= 5 for ODN, *n*= 5 HFD-saline, *n*= 5 for flumazenil + ODN.

To test whether endozepine signalling in the NTS is required for insulin regulation of HGP, we performed a pancreatic euglycemic-basal insulin clamp using flumazenil, a benzodiazepine antagonist, that prevents binding of DBI and ODN to the GABAA receptor. Surprisingly, flumazenil treatment prevented the insulin-dependent increase in GIR and decrease HGP (Figure 4B-Bii and S6), indicating endozepine modulation of GABAA receptors in the NTS is required for insulin signalling to the liver. Indeed, direct ODN infusion in the NTS could mimic the effect of insulin, causing an increase of GIR and a decrease of HGP (Figure 4B-Bii and S6). Neither flumazenil nor ODN had any effect on GU (Figure 4Bii). Together our data show that insulin acts directly on astrocytes (Fig. 2 & 4A) resulting in increased release of endozepines (Fig. 4 A). Moreover, we demonstrate that insulin sensing by NTS astrocytes is required for regulation of HGP (Fig. 3), which is prevented by blocking the action of endozepines on GABAA receptors (Fig 4B) and is replicated by increasing endozepines infused in the NTS (Fig. 4B). Interestingly, previous observations have shown that ICV injection of ODN improves glucose tolerance in mice^32^.

### Inhibition of GABAA receptors in the NTS decreases hepatic glucose production

Our data indicate that insulin likely modulates the activity of GABAA receptors by evoking release of endozepines from astrocytes. This provides a way for insulin to modulate neuronal activity in cells that lack the insulin receptor, for example, if the released endozepines reduce GABAA receptor function this would result in disinhibition of surrounding neurons. Such a scenario would be consistent with our data in Fig. 2, where insulin induced an increase of c-Fos in the NTS predominantly in neurons that did not uptake insulin. To determine whether a reduction in GABAA receptor activity in the NTS mimics the effect of insulin in reducing HGP, we used 2 different competitive GABAA antagonists (Fig. 5 and Supplementary Figure S4). Again, insulin injection in the NTS via the brain cannula, increased in GIR associated with a decrease in HGP while the glucose uptake was unchanged (Figure 5C-Cii and S7A-Ai). When bicuculine or gabazine was infused into the NTS it mimicked the effect of insulin, there was a clear increase in GIR and this was associated with a clear decrease in HGP without changes in GU (Figure 5A-Aii). These data suggest that the endozepines released from astrocytes are acting as negative modulators of NTS GABAA receptors. This is also the first demonstration that attenuating GABA signalling can reproduce the effect of insulin in the NTS.

**Figure 5.**
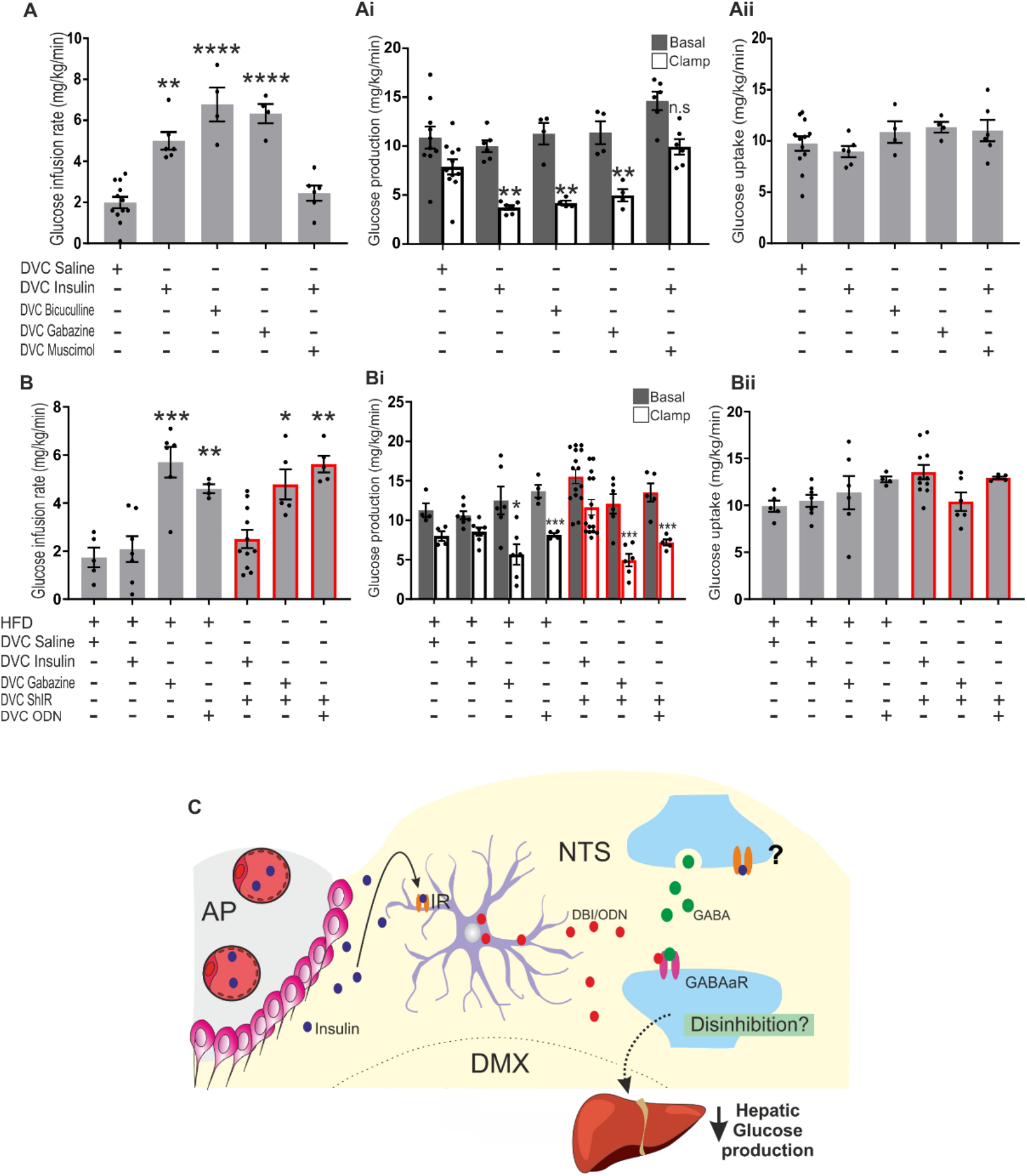
Effect of GABA inhibition/activation on insulin-dependent regulation of glucose production. Saline, insulin (a total of 2.52 mU, 1.26 mU per site), bicuculline ([1pM)) gabazine ([0.1mM]), or ODN ([10mM]) were infused at 0.006 ml/min during the clamp through the DVC catheter. Muscimol ([0.1µM])) was infused together with insulin 90 min after the experiment started. Where indicated ShIR was used to knock down the insulin receptor or **(A-B**) Glucose infusion rate (mg/Kg/min) during the final 30 min of the clamp for the indicated treatment. (**Ai-Bi**) Glucose production (mg/Kg/min) in basal (with square) and clamp (black square) conditions. (**Aii-Bii**) Glucose uptake was comparable in all groups. Values are shown as mean + SEM; *n*= 12 for saline, *n*= 5 insulin, *n*= 4 for Bicuculline and gabazine, *n*= 5 for Muscimol, *n*= 5 HFD-saline, *n*= 7 HFD-Insulin, *n* = 4 HFD-ODN, and *n*= 6 HFD-gabazine. Rats were fed with HFD 3 days before the camp to cause insulin resistance in B-Bii. (**C**) Schematic representation of working hypothesis

### GABAA receptor agonism prevents the insulin induced decrease in hepatic glucose production

Our data are consistent with a mechanism whereby insulin in the NTS decreases HGP by acting upon astrocytes, which in turn release endozepines to decrease GABAA receptor activity in neurons. To further test this model we used muscimol, a potent GABAA receptor agonist^33^ with the idea that directly activating GABAA receptors would prevent any insulin -> astrocyte-> endozepine induced decrease in GABAA activity. In this experiment insulin was infused for 90 min before the addition of muscimol and the start of the clamp (supplementary figure S4). Insulin alone was able to increase GIR and decrease HGP, but in the presence of muscimol, insulin action was completely blocked, we could see a clear decrease in GIR and increase in HGP while GU was not affected (Figure 5 A-Aii and S7Ai). Our data indicates that insulin action in the NTS requires negative modulation of GABAA receptors and we can prevent insulin-induced regulation of HGP by activating GABAA receptors in the NTS.

### Rescue of NTS control of HGP in insulin resistant models with gabazine and ODN infusion

HFD-feeding causes insulin resistance in the DVC and prevents insulin induced decreases in HGP. We hypothesised that using gabazine to inhibit downstream GABAergic signalling could be sufficient to bypass insulin resistance and decrease HGP. To this end we performed a basal glucose clamp study with the same protocol shown in Supplementary Fig. S4 where rats where fed with HFD for 3 days prior to performing the experiment. As expected, in HFD-fed rats, insulin injection in the NTS was not able to increase GIR or decrease HGP (Figure B to Bii and S7B). However, NTS-injection of gabazine was able to increase GIR by lowering HGP in HFD-fed rats (Figure 5B to Bii and S7B). Similarly, in HFD fed insulin resistant animals, infusion of ODN was also able increase GIR by lowering HGP (Figure 5B to Bii and S7B). These data suggest that the locus of the HFD-induced insulin resistance is likely astrocytic, as mimicking the downstream effects of insulin on astrocytes rescues the NTS’s ability to control HGP. Furthermore, selective knockdown of the insulin receptor in astrocytes recapitulates HFD-induced resistance (Figure 3G) and when we either supplied the endozepine ODN or inhibited GABAA receptor with gabazine we again could rescue the effect of insulin (Figure 5 B to Bii and S7Bi).

## Discussion

In summary, our data reveal that insulin action within the NTS leads to direct activation of approximately 20% of cells containing the insulin receptor (Fig. 2), yet widespread activation of cells lacking the insulin receptor. Notably, the majority of directly activated cells were found to be astrocytes. Through *in vivo* glucose clamp experiments, we demonstrated that knocking down the insulin receptor specifically in astrocytes effectively abolishes the NTS-mediated effect of insulin on regulation of HGP (Fig. 3G). Interestingly, astrocytes have been identified as capable of releasing endozepines ^23–25^ and we show that insulin stimulates the release of DBI in primary astrocytes (Fig. 4A). Furthermore, direct injection of ODN, the cleaved form of DBI, into the NTS is sufficient to decrease HGP (Fig. 4B). Both DBI and ODN are known to modulate GABAA receptors and we demonstrate that inhibiting binding of endozepines to the modulatory central benzodiazepine recognition site (CBR) on GABAA receptors blocks the effect of both insulin and ODN infusion in the NTS (Fig. 4). We also show that negative modulation of GABAA receptors is consistent with the effect of insulin and endozepines (Fig. 5A) whereas GABAA agonism prevents the effect of insulin (Fig. 5A). These data are consistent with astrocytic endozepines acting as negative allosteric modulators within the NTS. Similar negative allosteric actions of endozepines have been found in embryonic spinal cord neurones^34,35^, in the sub-ventricular zone^36^ and in CA1 pyramidal cells^37^. However, endozepines are also reported to have positive allosteric effects on the GABAA receptor in dorsal root ganglia^38^ and the dentate gyrus ^37^ indicating that endozepine action on GABAA receptors is not yet fully understood. It may depend on the subunit composition of GABAA receptors but clearly depends on the brain area, which is exemplified by Courtney *et al* (2018) who showed that within the hippocampus endozepines act as negative modulators within CA1 but positive modulators in the dentate gyrus ^37^.

When considering the receptors influenced by endozepines, there are three potential candidates. Firstly, there’s the GABAA receptor, responsive to the binding of both ODN and DBI. Secondly, ODN, accompanied by a smaller fragment (OP), can bind to a metabotropic G-protein-coupled receptor (GPCR) sensitive to PTX^26^. Additionally, endozepines also target the Translocator Protein (TSPO), capable of binding both ODN and DBI^26^. Through our use of flumazenil, we have definitively shown that the binding of DBI/ODN to the GABAA receptor is crucial for transducing insulin signalling in astrocytes of the NTS. Notably, studies conducted in the hypothalamus have suggested that ODN enhances POMC neuronal activity by acting on the metabotropic receptor. Indeed, OP can mimic the effects of ODN on regulating feeding behaviour and POMC activation. This effect is blocked by cyclo1-8[D-Leu5]OP, a competitive antagonist of ODN and OP^39^. Interestingly, ODN can also bind to the GABAA receptor and reduce inhibitory currents in POMC neurons, but this effect alone is insufficient to trigger activation of POMC neurons^39^. One could hypothesize that the regulation of glucose levels and feeding behaviour are associated with two different pathways. Perhaps the GABAA pathway is crucial for activating neurons that signal via the vagus to the liver, while the metabotropic pathway is used for activation of POMC neurons to control feeding. Indeed, it has been shown that in the NTS, POMC neurons are activated by OP treatment, and this leads to a decrease in food intake^40^.

An extensive body of work has shown that endozepines can modulate feeding behaviour by acting both on the hypothalamus and in the brain stem ^41–46^. Our findings reveal that insulin stimulation prompts the release of DBI from primary astrocytes, and the subsequent administration of ODN directly into the NTS effectively reduces HGP. This is the first instance of establishing a direct link between insulin signalling and the astrocyte-dependent release of endozepines. Before our study, only limited evidence existed regarding the role of brain-derived endozepines in controlling glycemia. Notably, a study in 2013 demonstrated that injecting ODN into the hypothalamus could improve glucose tolerance, though the precise mechanism remained unclear^32^. Remarkably, intraperitoneal injection of insulin elicits an upregulation of DBI mRNA expression in the ventromedial hypothalamus ^47^, and glucose infusion in fasted rats leads to an elevation in DBI mRNA levels within the hypothalamus ^32^. Conversely, fasting has been shown to decrease DBI mRNA levels in the hypothalamus ^46^. These data suggest that, changes of energy status that lead to elevated circulating insulin levels can favour the increase of DBI transcription. ^47^, and our study furthers these observations by demonstrating the insulin evokes release of DBI from astrocytes within the NTS.

Although our study has focused on the effect of insulin receptors in astrocytes, it is worth noting that the insulin receptor is also present in neurons, as illustrated in Figure 1. Whether a direct action of insulin on neurons plays a significant role in regulating HGP remains unclear. There is a clear body of evidence, including our previous work that suggest that insulin activates the KATP channel to decrease HPG^9,10^. Intriguingly we have shown that the majority of the neurons that uptake insulin are not activated (Fig. 2E), consistent with the hyperpolarising effect of activating KATP channels. Perhaps, across the population of NTS neurons, insulin acts as a toggle switch, dialling down activity in neurons containing the insulin receptor and KATP channels, while boosting activity in other neurons by dialling down the effects of GABAergic inhibition with astrocytic release of DBI/ODN. Further investigation should aim to elucidate the effects of insulin and DBI/ODN on NTS neuronal subtypes.

Short term high fat diet (HFD)-feeding in rodents causes insulin resistance and impairs the ability of the DVC and mediobasal hypothalamus (MBH) to respond to insulin preventing regulation of HGP and food intake^10,11,48^. Interestingly, in both the MBH and the DVC, HFD causes alteration of mitochondrial dynamics and increases in ER-stress, these events in turn lead to loss of insulin sensitivity and alter the ability of insulin to regulate blood glucose levels and feeding behaviour ^20,49–51^. It is sufficient to inhibit mitochondrial fragmentations in NTS-astrocytes to prevent the development of insulin resistance, thus suggesting that astrocytes are key insulin sensors in the NTS. Indeed, here we have shown that in HFD-fed insulin-resistant rats or in rats where the insulin receptor has been knocked out in the NTS, mimicking the downstream effects of astrocytes by infusion of ODN or gabazine (Fig. 5B) can bypass the insulin resistance and decrease HGP.

In summary, our research is beginning to unravel the intricate molecular and cellular events underlying insulin’s action in the NTS-mediated modulation of glucose levels. Astrocytes have been identified as the primary site of insulin action, and the release of endozepines appears to play a crucial role in modulating GABAergic signalling to regulate glucose production (Fig. 5C). Further investigation is warranted to elucidate the precise mechanisms and contribution of insulin-dependent astrocyte-neuronal crosstalk among different neuronal populations within the NTS.

## Materials and Methods

### Animal Preparation

Our studies utilized nine-week-old male Sprague-Dawley rats, weighing between 280 and 320 g, obtained from Charles River Laboratories, (Margate, UK). Animals were used in line with the United Kingdom animals (Scientific Procedures) Act 1986 and ethical standards set by the University of Leeds Ethical Review Committee. Animals were housed individually in cages and maintained under a standard 12-hour light-dark cycle with ad libitum access to standard rat chow or high fat diet and water. On day 0 rats underwent stereotactic surgery (World Precision Instruments)) for implantation of bilateral catheters (Plastics One, Virginia, USA) targeting the nucleus of the solitary tract (NTS) within the dorsal vagal complex (DVC) (coordinates: 0.0 mm on occipital crest, 0.4 mm lateral to midline, and 7.9 mm below the skull surface)^10^

For rats in the GFAP-ShIR cohort, on the day of brain surgery, rats were co-injected with an adenovirus expressing the Cre recombinase under the GFAP promoter (GFAP-Cre) and with an adenovirus expressing a FLEx plasmid where ShRNA for the insulin receptor and tdTomato (ShIR) are cloned within 2 LoxP sites. A scrambled version of the ShRNA was used as a control (ShC)

For animals undergoing pancreatic-euglycemic clamp, following a one-week recovery period, the rats underwent intravenous catheterization, with catheters inserted into the internal jugular vein and carotid artery for infusion and sampling purposes (figure S4).

### Virus preparation

To inhibit the insulin receptor we utilized a Cre-LoxP system where pAAV[FLEXon]- CMV>LL:rev(tdTomato:rInsr[miR30-hRNA#3]):rev(LL):WPRE and pAAV[FLEXon]- CMV>LL:rev(tdTomato:Scramble[miR30-shRNA#1]):rev(LL):WPRE control scramble vectors were designed and purchased from Vector Builder. ShRNA (short hairpin RNA) was designed to silence target mRNA of insulin receptor (ShIR), and cloned into an adenoviral shuttle vector to make pacAd5CMV-ShIR-tdTomato and pacAd5CMV- ShC-tdTomato, the scrambled RNA possesses the same nucleotide composition as the ShIR but is incongruent with any target mRNA of IR. This was used as control vector and named as ShC. In addition, a pacAd5CMV-CRECFP vector was designed and generated to introduce Cre recombinase. To produce an adenoviral system expressing the Cre under the GFAP promoter, the CMV promoter was removed from the pacAD5 shuttle vector, and replaced with the rat GFAP promoter. Recombinant adenoviruses were amplified in HEK 293 AD cells and purification was performed using a sucrose-based method^20^. Adenoviruses injected into the animals’ brains had a titre between 3 × 109 pfu/ml and 4.4 × 1011 pfu/ml

### In vivo FITC-insulin DVC infusion

Animals were fitted with a bilateral DVC cannula as detailed previously and were allowed to recover for 1 week. Following recovery, animals were injected with 495uM FITC-insulin (2uL per site) into the NTS, the cannula was left in place for 5 minutes before being removed and replaced with a dummy cannula. After 46 minutes animals were terminally anaesthetized and perfused with 4% PFA, brains were dissected and prepared for either RNAscope or immunofluorescence (figure S1) as detailed below.

### RNAScope and Immunofluorescence

Animals were terminally anaesthetized (Pentoject, 60 mg/kg IP) and transcardially perfused with 4% paraformaldehyde. The brainstem area containing the DVC was taken and frozen in optimal cutting temperature compound (OCT, VWR international); 10 μm and 25 μm sections were cut using a cryostat for RNAScope and immunofluorescence, respectively.

FITC-insulin+ sections were permeabilised in 0.3% PBST (1X PBS, 0.3% Triton X-100 (PBST)) for 10 minutes and blocked for 1 hour with 10% normal donkey serum (Merck Millipore), 1% bovine serum albumin (BSA) (Sigma Aldrich) in 0.3 M glycine (Sigma Aldrich) PBS to prevent nonspecific background labelling. Sections were then incubated c-Fos (1:400 in 0.1% PBST, Cell Signaling Technology, #2250) overnight at 4°C. After 24 hours sections were washed in PBS and then incubated with Alexa Fluor® 555 donkey anti-rabbit (1:1000 in PBS, Thermofisher, #A32794) for 2 hours at room temperature, Following PBS wash, sections were mounted using Vectashield plus DAPI (Vector laboratories, H-1200-10).

RNAScope Multiplex Assay V2 (Bio-Techne) was performed according to manufacturer’s instructions in order to characterise the expression of INSR (probe cat# 406421) or FITC-insulin with or without c-Fos (cat# 403591) in either GFAP+ (cat# 407881-C2) astrocytes, VGAT+ (cat# 424541-C3) inhibitory neurones, or VGLUT+ (cat# 317011-C2) excitatory neurones. Sections were stained with DAPI to label nuclei and imaged using either Zen AxioScan V1 slidescanner or LSM880 upright confocal microscope.

Images were analysed using the cell counter plug in in Fiji (Image J). Cells positive for INSR plus CFOS, VGAT, VGLUT, or GFAP were counted in 3 regions of interest (ROI) from the NTS from 3 sections containing the DVC from 3 animals, giving a total of 9 regions of interest per section per animal to give a total of 27 ROIs analysed.

To determine the specificity of the GFAP-ShIR-tdTomato expression sections were checked for the presence of tdTomato in the DVC and co-labelled with GFAP (1:1000 in 0.1% PBST, Abcam ab7260) at 4°C overnight. Following incubation with appropriate secondary antibody (Thermofisher, #A32794) sections were mounted using vectashield plus DAPI. Images were taken using a Zeiss LSM880 upright confocal laser scanning microscope.

### DVC Infusion and Pancreatic-Euglycemic Clamp Studies

Four days following intravenous catheterization, animals whose food intake and body weight had returned to within 10% of baseline levels underwent the clamp studies (figure S4). To ensure consistent nutritional status during the clamps, rats were limited to 15 g (50.7 cal for RC and 77.1 cal for HFD) of food the night before the experiment. The clamps lasted for 210 minutes. During the clamp period (from time 0 to 210), saline, insulin (totaling 1.26 μU per site) bicuculline ([1pM]), gabazine ([0.1mM]), or ODN ([10mM]) were infused at 0.006 ml/min during the clamp through the DVC catheter. Muscimol ([0.1µM])) was infused together with insulin 90 min after the experiment started. For experiments using the CBR site antagonist flumanzeil, flumazenil ([33nM]) was pre-infused at 0.006 μl/min for 1 hour and then co-infused with insulin or ODN during the clamp Additionally, a primed-continuous intravenous infusion of [3-3H]glucose (40 μCi bolus, 0.4 μCi/min) was initiated at 0 minutes and maintained throughout the study to assess glucose kinetics. A pancreatic (basal insulin)-euglycemic clamp was initiated at 90 minutes and continued until 210 minutes. Intravenous infusion of somatostatin (3 μg/kg/min) along with insulin (0.7-1 mU/kg/min) was performed to return insulin levels to basal levels. For HFD fed animals insulin was infused at 1.4-1.8mU/kg/min). A 25% glucose solution was administered starting at 90 minutes and periodically adjusted to maintain plasma glucose levels. Samples for the determination of [3-3H]glucose-specific activity and insulin levels were collected at 10-minute intervals.

### Biochemical Analysis

Plasma glucose concentrations were measured by the glucose oxidase method (Glucose Analyzer GM9; Analox Instruments, Stourbridge, UK). Plasma insulin levels were determined by an ultra-sensitive rat insulin ELISA low range assay (0.1–6.4 ng/ml) (Crystal Chem, Elk Grove, IL, USA) used according to manufacturer’s instructions

### PC12 cell culture, insulin receptor knockdown, and FITC-insulin treatment

Pheochromocytoma (PC12) cells (AddexBio # C0032002) were cultured in F-12K medium (Gibco # 21127-022) with 15% horse serum (Gibco # 1011-07), 2.5% fetal bovine serum (Gibco # 2024-01), and 1% Penicillin Streptavidin (Sigma # P0781) under 37 °C, 5% CO2 conditions. A confluent 75 mm flask of PC12 cells was diluted 1:3 and plated in a 10 cm dish containing a multi-test 8-well slide (MP Biomedicals # 20190128). 24 hours later cells were transfected with 5 μg pacAd5CMV-CRECFP and

5 μg pacAd5CMV-ShIR-tdTomato or 5 μg pacAd5CMV-ShC-tdTomato control by using GenJet reagent (SL00489-PC12). Fresh complete media was added to the cells after 24 hours. 48 hours later, cells were incubated with 15 uL of 800nM FITC-insulin for 15 minutes before being fixed with 4% PFA and then washed with 1 X PBS 3 times. Excess liquid was then aspirated, and the slide left to dry. Cover slips were added and sealed with Vectashield mounting medium plus 4’,6-diamindino-2-phenylindole (DAPI) to visualise nuclei (Vector Laboratories, Burlingame CA, USA). Slides were imaged with an upright confocal microscope (Zeiss LSM880)

### Primary astrocyte culture, insulin receptor knockdown, and insulin treatment

Rat pups (postnatal day 1-3) were decapitated and the cortices were dissected and centrifuged at 1000rpm for 5 minutes. The supernatant was removed, 2ml of trypsin was added and left to incubate for 30 minutes. 2mg/ml DNase was added and samples were centrifuged for 5 minutes at 1000g. Once the supernatant was removed 10ml of culture media (10% FBS, DMEM, 1% PSF) was added and the cells were suspended by repeated pipetting and placed in a Poly-D-lysine coated T75 flask. The following day media was changed and 10 uL PLX was added. One week later, when cells were confluent, flasks were placed on an orbital shaker at 180rpm for 30 minutes to detach microglia. The microglia containing media was removed and replaced with fresh media and flasks were left on an orbital shaker at 240 rpm for 6 hours to remove oligodendrocyte precursor cells. The culture media was removed and flasks were rinsed with PBS twice and trypsin added to detach the astrocytes. Astrocytes were pelleted and then resuspended in 40ml of fresh astrocyte medium and plated into a Poly-D-lysine coated T75 flask.

For insulin receptor knock down 3-4 days prior to insulin treatment primary astrocytes where infected with 2 adenoviruses expressing Cre-recombinase and a FLEx virus expressing the ShIR or the ShC. Cells where then starved for 5 hours and treated with 100nM insulin or saline for 30 min. Cells where then harvested, lysed and analysed by western blotting, and probed with the indicated antibodies as detailed below.

### Western Blotting

PC12 or primary astrocyte cells were lysed in lysis buffer (50 mM Tris-Hcl pH 7.5, 1mM EGTA, 1mM EDTA, 1% (w/v) NP-40, 1mM sodium orthovanadate, 50 mM sodium fluoride, 5mM sodium pyrophosphate, 0.27M sucrose, 1μM DTT and Pierce Protease Inhibitor Tablets), using a homogeniser on ice. Samples were centrifuged at 12,000 RPM for 15 min at 4° supernatants were collected and protein concentration was determined by using a Bradford Assay. Proteins were separated and measured by western blotting as previously described^20^. Membranes were blocked in 5% BSA TBST and incubated in the primary antbodies anti-INSR (Cell Signaling Technology, #3025), anti-TdTomato (SICGEN, AB8181-200), anti-p-AKT (Cell Signaling Technology, #9271) anti-T-AKT (Cell Signaling Technology, #9272S), anti-β-actin (Cell Signaling Technology, #3700) overnight at 4°C, followed by incubation with species appropriate secondary HRP-conjugated antibodies (Invitrogen, Life Technologies) in 5% skimmed milk. Membranes were imaged using ECL (BioRad Clarity) with the iBright FL1500 Imaging system and analysed using iBright Analysis software (Thermofisher).

### Statistical analysis

All data are presented as mean ± SEM and where applicable data from each animal is reported as a single data point. Analysis was performed using GraphPad Prism 9 software. Statistical significance was assessed using multiple T-tests or one-way ANOVA (with Sidak’s post-hoc test). “*N*” denotes the number of animals used. A p-value less than 0.05 was considered statistically significant. Levels of significance were categorized as follows: (*) p < 0.05; () p < 0.01; (*) p < 0.001; (****) p < 0.0001.

## Acknowledgments

This work was supported by an MRC-Career Development Fellowship (MR/S007288/1) that also provided salary BMF and LEN and by a Wellcome Trust (UNS63234) grant. SK was funded by the European foundation n for the study of Diabetes (EFSD)-Multi-System Challenges in Diabetes” 2021. Microscopy was performed in the bioimaging facility in Leeds and the Zeiss LSM 880 inverted confocal was funded by Wellcome Trust (104918/Z/14/Z).

## Author contributions

**L.E.N.** Conducted and designed experiments, performed data analyses and wrote part of the manuscript. **N.W** Conducted FITC-insulin experiments in vitro. **S.K.** Conducted experiments in primary astrocytes. **J.C.G.** Assisted with the experiments and produced the adenoviruses. **R.H**. Performed the analysis in Figure 2. **J.J.** participated in experimental design and data interpretation. **B.M.F.** Conceived and supervised the project, designed the experiments (and conducted some), and wrote the manuscript.

## Declaration of interests

No conflict of interest

## Supplementary material

**Figure S1:**
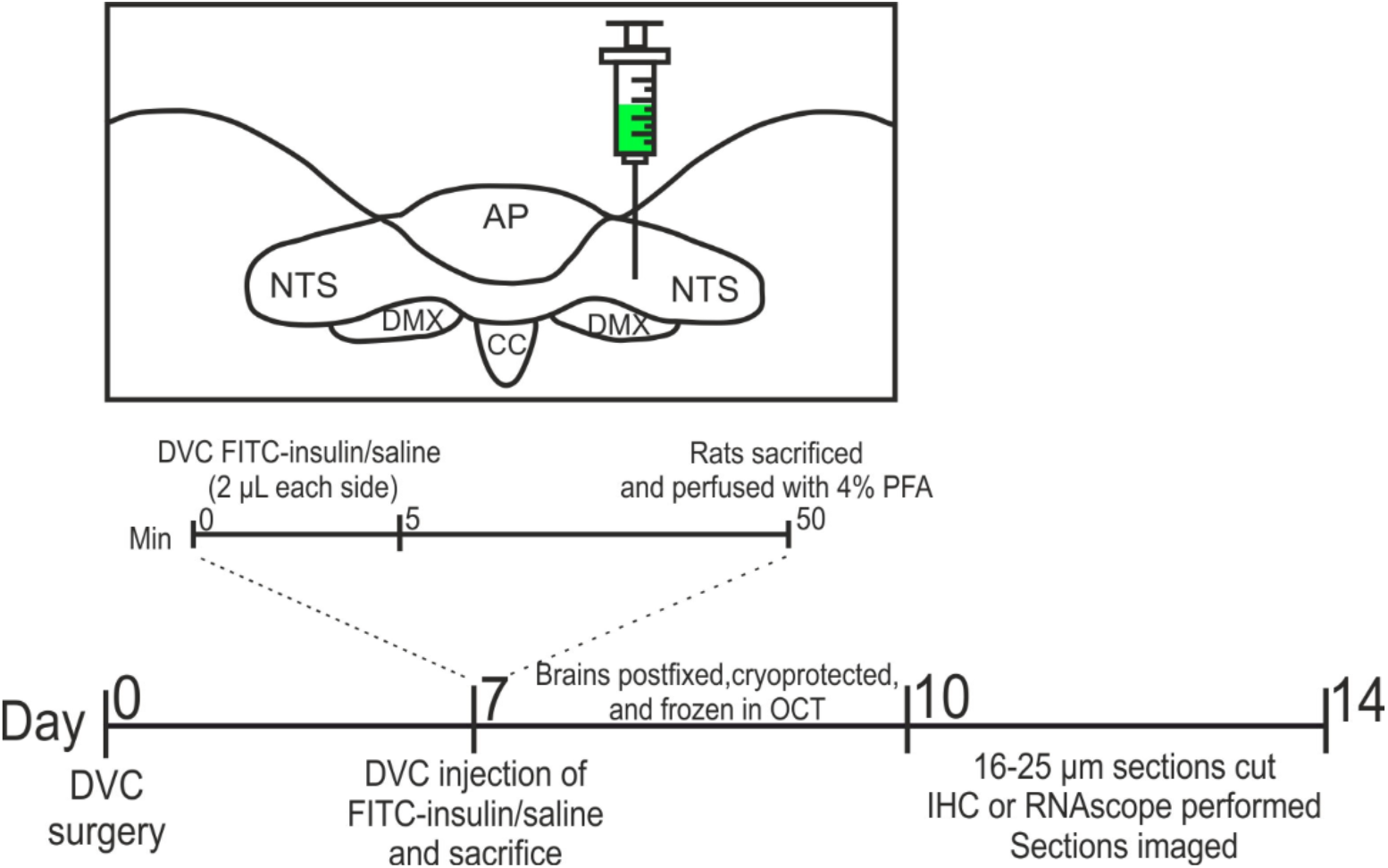
Schematic of FITC-insulin NTS injection paradigm related to figure 2.

**Figure S2:**
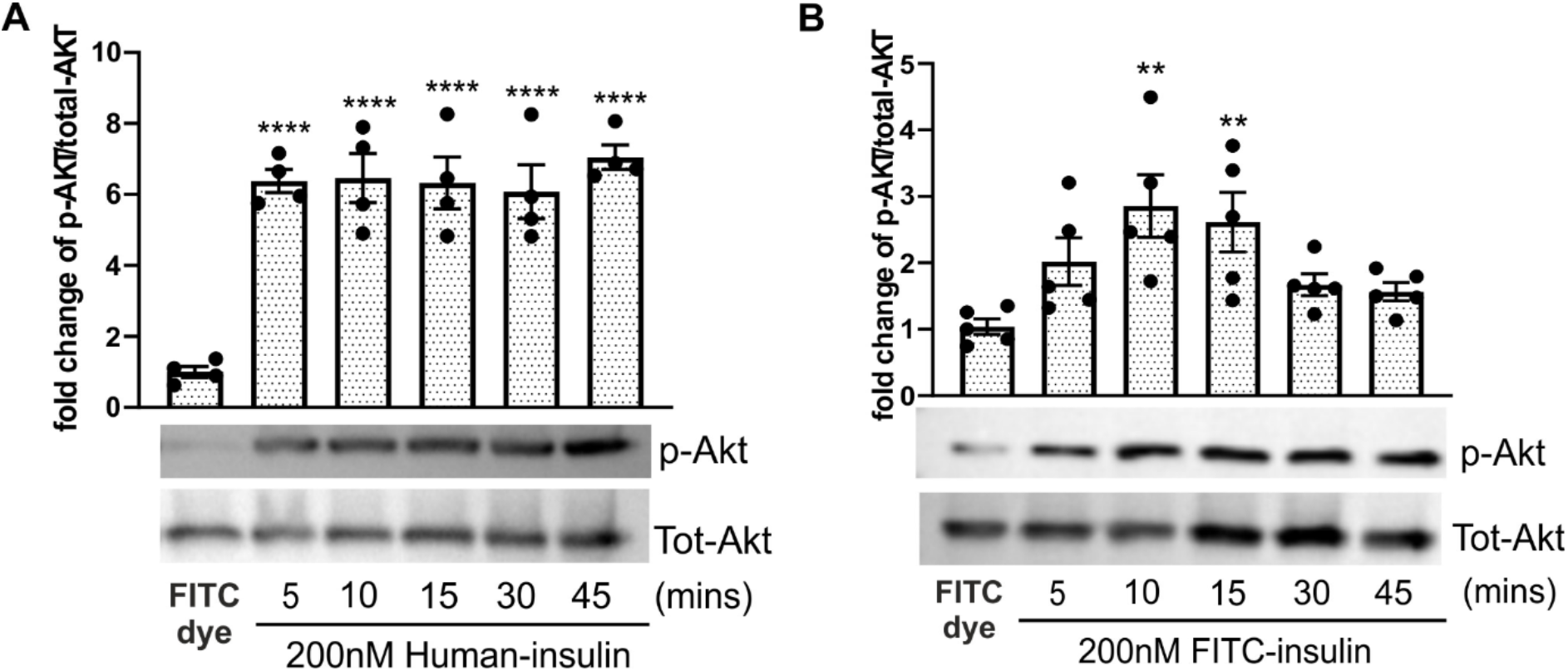
Comparison of the effect of human insulin and FITC-insulin treatment on AKT activation in PC12 cells.

**Figure S3:**
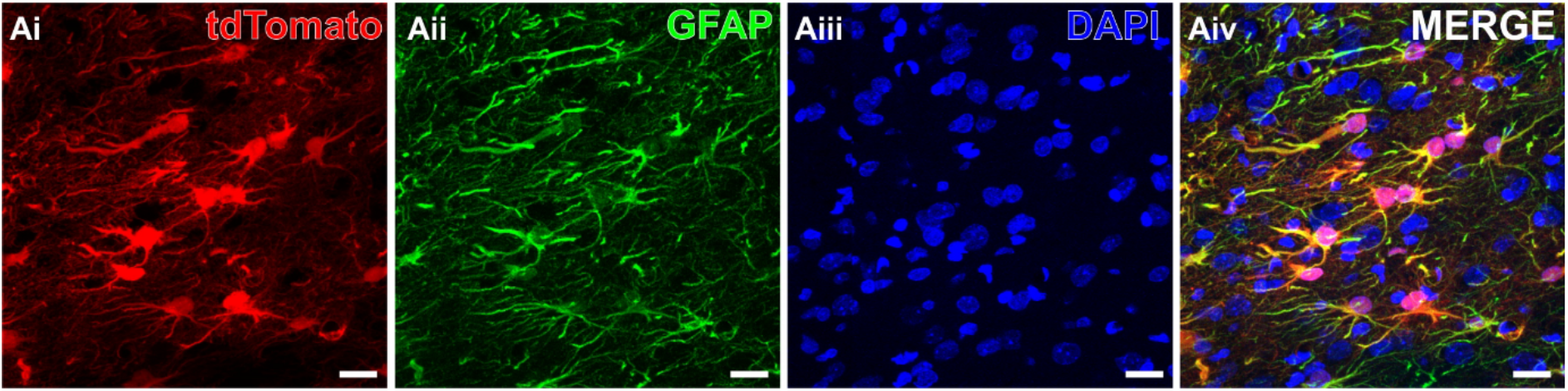
Representative confocal image of expression of tdTomato+ ShIR in GFAP+ astrocytes in the NTS – related to figure 2.

**Figure S4:**
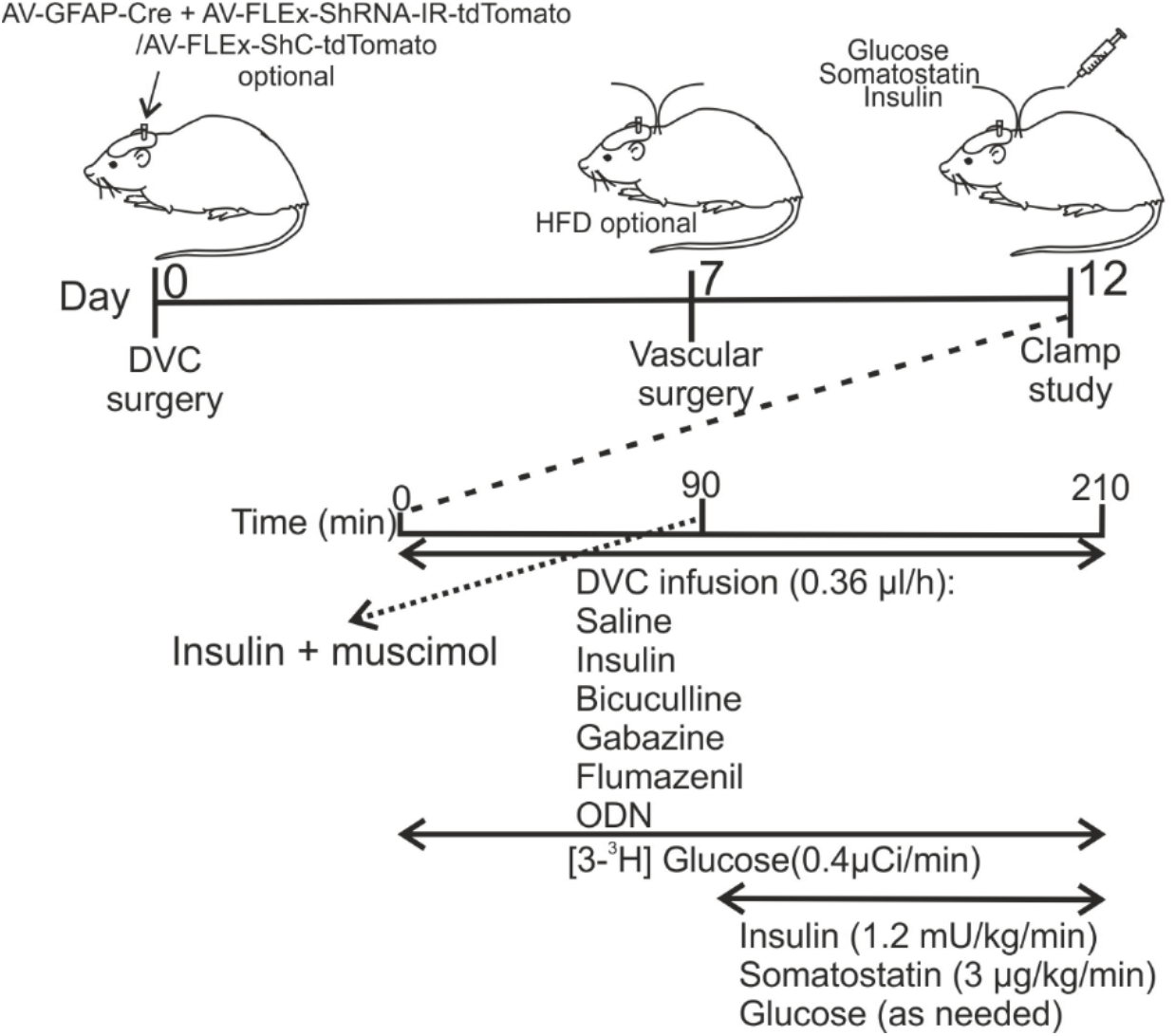
Schematic of pancreatic- euglycemic clamp protocol.

**Figure S5:**
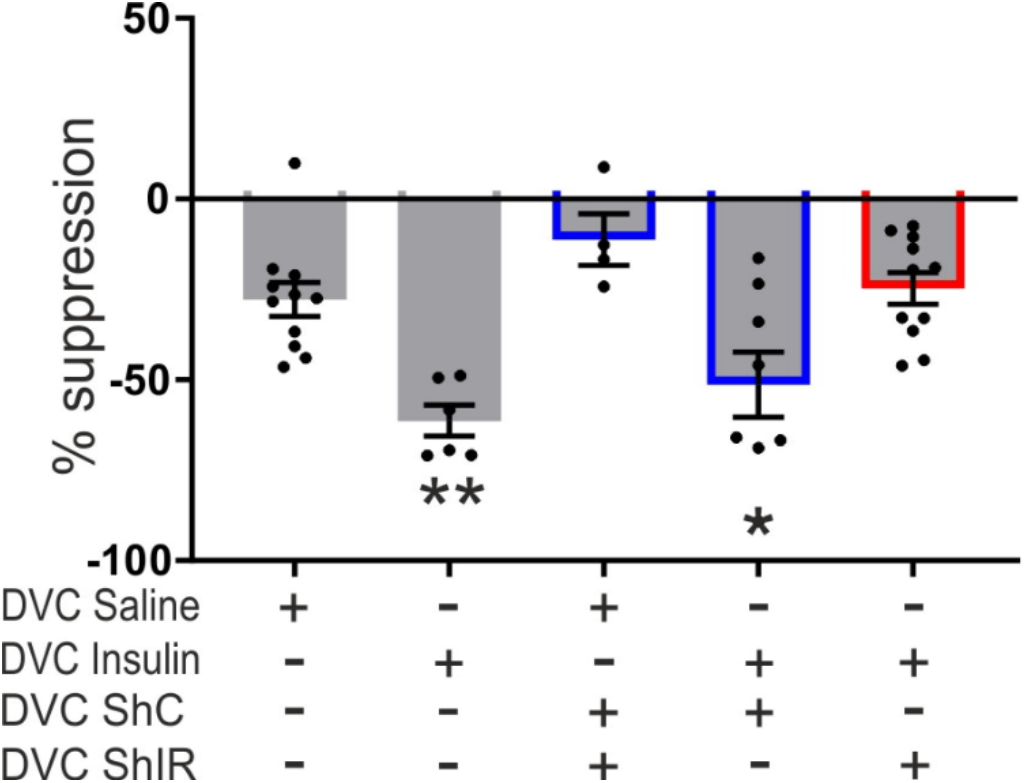
Glucose kinetics during the clamps with DVC saline or insulin treatment in animals expressing either ShC or ShIR in NTS GFAP+ astrocytes related to figure 3.

**Figure S6:**
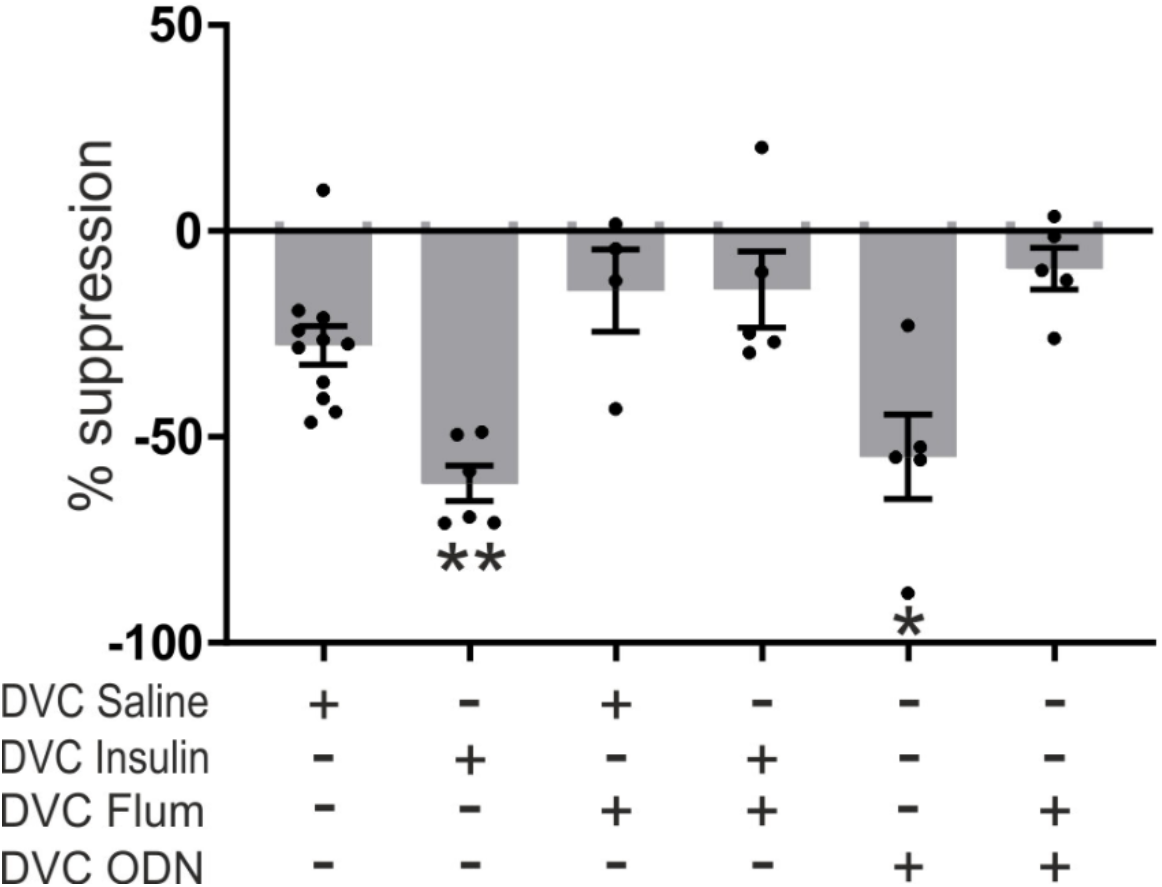
Glucose kinetics during the clamps with DVC flumazenil or ODN treatment in animals related to figure 4.

**Figure S7:**
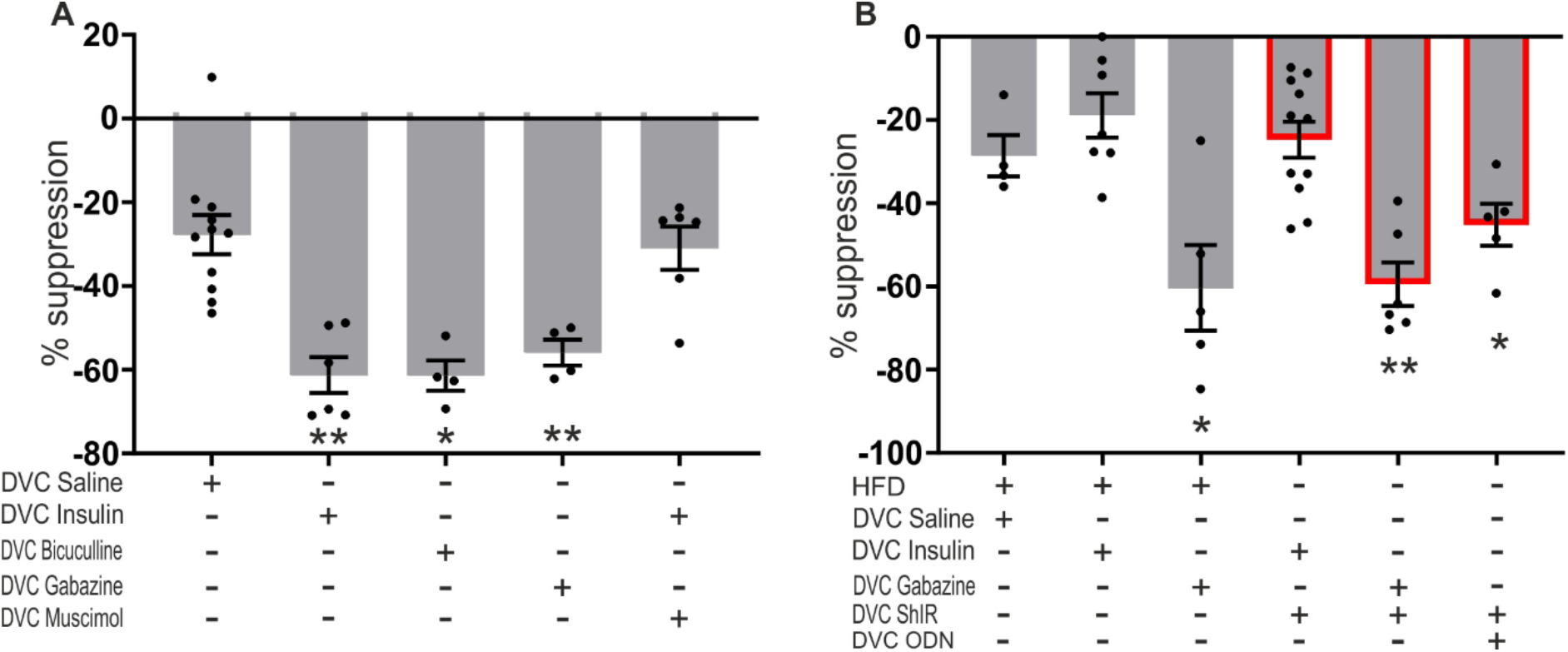
Glucose kinetics during the clamps in RC fed animals treated with either DVC bicuculline, gabazine, muscimol treatment (A-Ai). (B-Bi) Glucose kinetics for either HFD fed or GFAP-ShIR-tdTomato+ animals treated with either gabazine, flumazenil, or ODN related to figure 5.

**Table S1:**
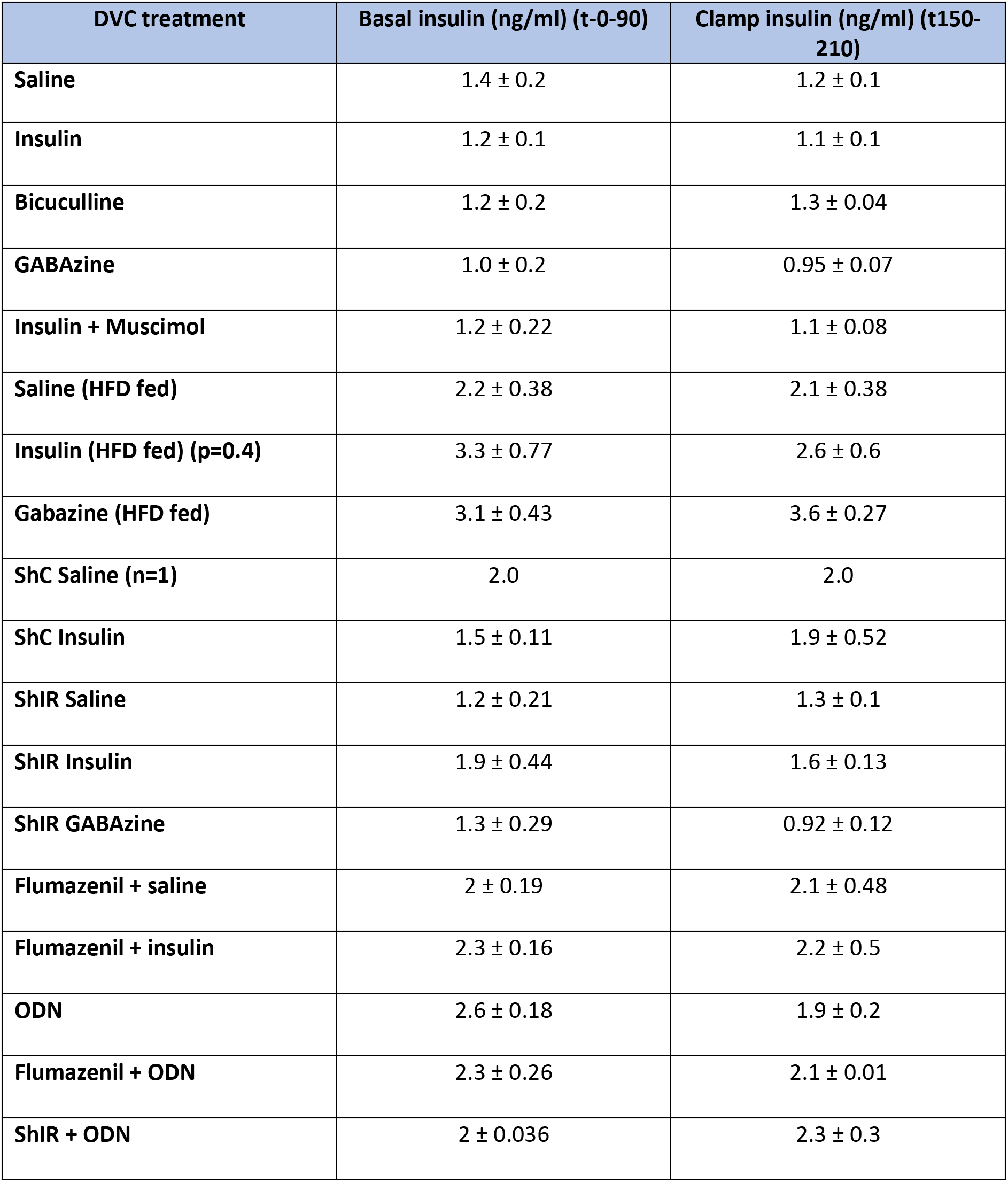
Summary of basal and clamp insulin levels (ng/ml) during the pancreatic- euglycemic clamp experiments.

**Table S2:**
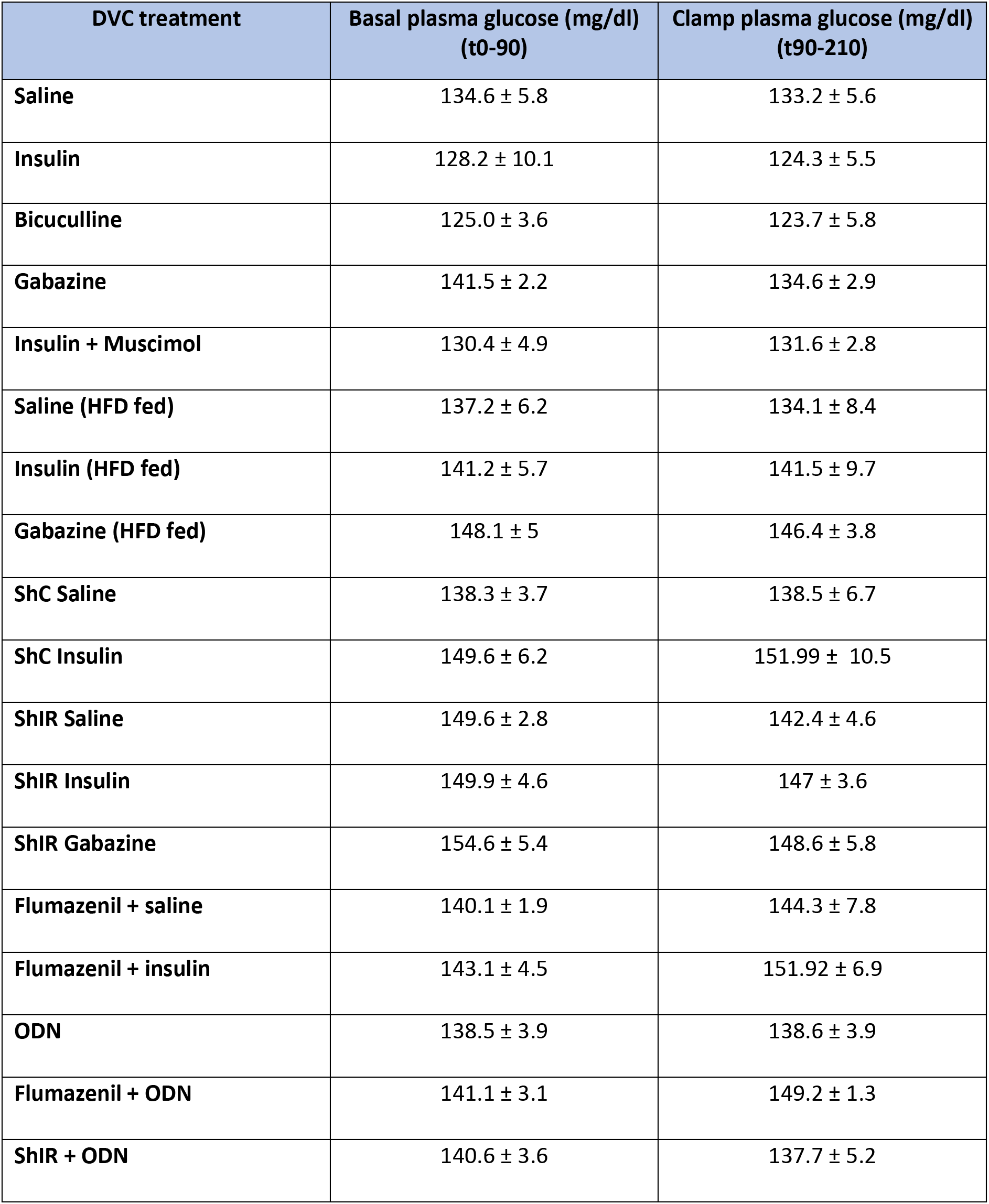
Summary of basal and clamp plasma glucose levels (mg/dl) during the pancreatic- euglycemic clamp experiments.

## Notes

### Competing Interest Statement

The authors have declared no competing interest.

